# Cytoneme-mediated transport of active Wnt5b/Ror2 complexes in zebrafish gastrulation

**DOI:** 10.1101/2022.04.07.487468

**Authors:** Chengting Zhang, Lucy Brunt, Sally Rogers, Steffen Scholpp

## Abstract

Chemical signaling is the primary means by which cells communicate in the embryo. The underlying principle refers to a group of ligand-producing cells and a group of cells that respond to this signal because they express the appropriate receptors. In the zebrafish embryo, Wnt5b binds to the receptor Ror2 to trigger the Wnt/Planar Cell Polarity (Wnt/PCP) signaling pathway to regulate tissue polarity and cell migration. However, it is still unclear how this lipophilic ligand is transported from the source cells through the aqueous extracellular space to the target tissue. Here we show that Wnt5b, together with Ror2, is loaded on long protrusions. The active Wnt5b/Ror2 complexes are handed over from these cytonemes to the receiving cell to trigger Wnt/PCP signaling, regardless of whether the cell expresses functional receptors. On the tissue level, we show that cytoneme-dependent spreading of active Wnt5b/Ror2 affects convergence and extension in the zebrafish gastrula.

## Introduction

Chemical signals are one of the main modes of intercellular communication in embryonic development. The signal is released by signal-producing cells in the form of secreted molecules called ligands. The target cells have the ‘ability to react’ and respond to this signal by changing the cellular behavior (Spemann and Mangold, 1924). The ability to transduce such a signal depends, foremost, on the availability of the correct set of receptors. Consequently, cells and tissues lacking the appropriate receptors are considered non-responsive or blind. Signal activation in responsive cells but not in the non-responsive cells allows for precise tissue specification, organ development, and formation of the entire embryonic body (Niehrs, 2004).

During embryonic development, one of the most critical signaling systems is the Wnt signaling network, which comprises evolutionarily conserved and entangled pathways. The Wnt signaling network regulates multiple processes crucial to embryogenesis and tissue homeostasis in all metazoans. The Wnt/Planar Cell Polarity (PCP) signaling is one pathway of this network and regulates cell polarization and migration (Tada and Heisenberg, 2012; Yang and Mlodzik, 2015). In vertebrates, crucial components of the Wnt/PCP pathway include the Wnt5 ligands, Wnt5a and Wnt5b, and the cognate tyrosine kinase-like orphan receptor 2 (Ror2) (Rogers and Scholpp, 2021; Stricker et al., 2017). In zebrafish, Wnt5/Ror2 regulates c-Jun N-terminal kinase (JNK), Rac, and RhoA signaling (Nishita et al., 2010; Oishi et al., 2003), which governs the concomitant narrowing and lengthening of the embryonic body axis, the so-called convergence and extension (C&E) movement (Roszko et al., 2009; Schlessinger et al., 2009; Wallingford et al., 2002). However, it is still unclear how the lipid-modified and membrane-associated Wnt5 ligands, produced at the embryonic margin, are transported over several hundreds of micrometers to cells of the overlying epiblast expressing their receptors to activate paracrine PCP signaling and thus control C&E in the embryo.

Cytonemes are thin and actin-rich membranous protrusions that transport essential components of signaling pathways between cells (Roy and Kornberg, 2015; Zhang and Scholpp, 2019). Cytonemes are highly dynamic and can form and retract within minutes. Their emergence is precisely controlled by the cytoneme-producing cell and the extracellular space they traverse. Cytonemes have been reported to transport signaling components of the Wnt signaling family in invertebrates and vertebrates (Zhang and Scholpp, 2019). For example, cytonemes can transport Wnt8a over tens of micrometers in the zebrafish gastrula to regulate the patterning of the neural plate (Stanganello et al., 2015).

Similarly, Wnt3 is loaded on cytonemes in gastric cancer to facilitate tumor proliferation (Routledge et al., 2022). The emergence of Wnt cytonemes is controlled by Wnt/PCP signaling in the source cell (Brunt et al., 2021; Mattes et al., 2018). However, it is unclear how Wnt ligands loaded on cytoneme are handed over to the receiving cells to engage with their cognate receptors.

Here, we explore the dissemination of the Wnt/PCP signaling components, Wnt5b, and its receptor Ror2 in zebrafish gastrulation. First, we show that fibroblasts and epiblast cells in the zebrafish embryo form cytonemes decorated with Wnt5b and Ror2. Our studies further show that these form a complex and are handed over together to the receiving cell. Then, using *in vivo* quantitative imaging with single-molecule sensitivity, we demonstrate that the cohesiveness of the Wnt5b/Ror2 complex is maintained in the producing cells, during transport along cytonemes, and in the receiving cell. Surprisingly, the conveyed complex remains active during the transport and regulates PCP signaling in the target cell, even when these cells lack functional Ror2 receptors. We further show that blockage of Wnt/PCP cytonemes influences the C&E movement in the zebrafish embryo. In summary, we show that cytoneme-mediated transfer of the Wnt/PCP ligand-receptor complex is vital for paracrine Wnt/PCP signaling and thus challenges the categorization of tissues into responsive and non-responsive on the receptor level.

## Results

### Cellular protrusions can disseminate wnt5b/Ror2 complexes

Wnt5 is one of the primary regulators of Wnt/PCP signaling in vertebrate embryonic development (Rogers and Scholpp, 2021). However, it is still unclear how these lipid-modified ligands are precisely disseminated in an embryonic tissue to regulate complex tissue movements like C&E. Therefore, we asked how Wnt5 spreads between cells. First, we investigated the localization of the ligand in zebrafish fibroblast (PAC2) by using our combined filopodia preservation and immunohistochemistry protocol (Rogers and Scholpp, 2020). By using an antibody against Wnt5a/b, we found that the ligand can be detected on over 65% of protrusions of PAC2 cells (Fig. 1A, B; Suppl. Fig. 1A). The Wnt5a/b positive protrusions are significantly longer than protrusions on which we cannot detect a signal. Next, we addressed the localization of Ror2, the cognate receptor of Wnt5a/b (Stricker et al., 2017), and we noticed Ror2 on cellular extension (Fig. 1C, Suppl. Fig. 1B). In addition, we find a robust localization of Ror2 in the nucleus as documented in other cell lines (Carbone et al., 2018). We then asked if the ligand and the receptor form clusters at the membrane. We found co-localization of the fluorescent-tagged ligand Wnt5b with the endogenous receptor protein. Vice versa, we detected co-localization of the fluorescent-tagged receptor Ror2 with endogenous Wnt5a/b protein - specifically on cellular extensions (Fig. 1D). These findings suggested that Wnt5a/b and Ror2 can co-localize on protrusions. To document the dissemination of these signaling components, we overexpressed the tagged constructs of the ligand Wnt5b and the receptor Ror2, individually and in combination in PAC2 fibroblasts (Fig. 1E). We found Wnt5b-GFP and Ror2-mCherry on protrusions. In parallel, we also detect co-localization of Wnt5b and Ror2 in puncta in the neighboring cells, suggesting that Wnt5b together with Ror2 can be handed over to adjacent cells. To investigate if endogenous Ror2 can also be transported to neighboring cells, we generated a Ror2 knock-out PAC2 cell line by CRISPR/Cas9 (Suppl. Fig. 1C, D). We co-cultivated the KO cells with WT PAC2 cells and stained these cells with a Ror2 antibody combined with Phalloidin to mark the actin cytoskeletal elements. (Fig. 1F). We used the prominent nuclear localization of Ror2 (Carbone et al., 2018) to distinguish between Ror2^-/-^ cells from Ror2 WT cells. We found that Ror2 is localized in clusters at the plasma membrane and in the nuclei in the WT cells (Fig. 1 F, G). By high-resolution imaging, we could identify Ror2 clusters in Ror2^-/-^ cells (Fig. 1G), suggesting that Ror2 protein was produced in the WT cells and then transferred to the KO cells.

**Figure 1.**
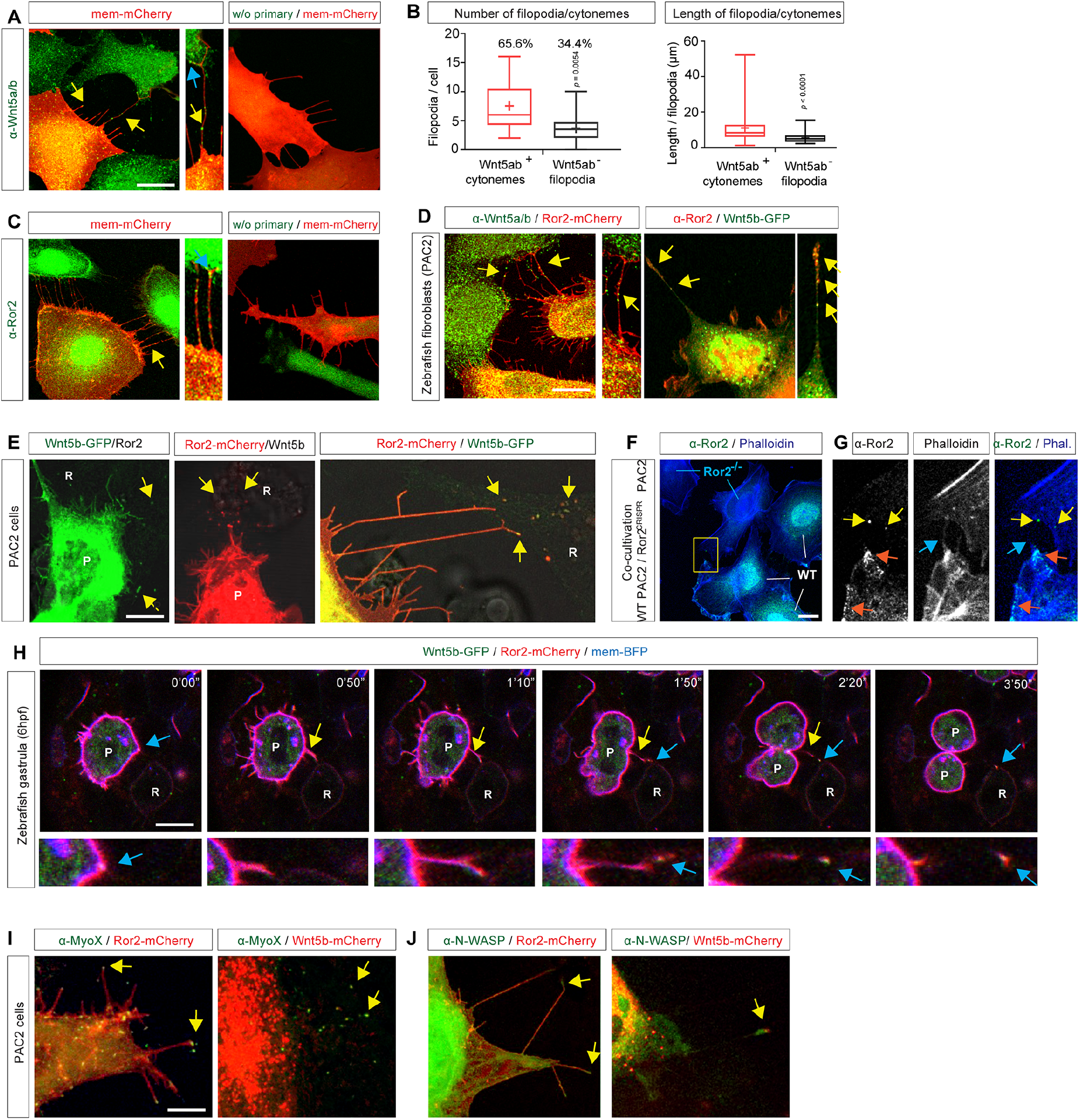
Wnt5b/Ror2 complexes are transported from the producing cells to receiving cells via cytonemes. **A**. PAC2 zebrafish fibroblasts were transfected with memCherry and stained with an antibody against Wnt5a/b. A high-magnification image indicates Wnt5a/b-bearing cytonemes. Yellow arrows indicate Wnt5a/b clusters on cytonemes. Scale bar 10µm. **B**. Quantification of the number and length of filopodia versus Wnt5a/b cytonemes of PAC2 cells (n=17, n= numbers of cells). *p*-values were determined by a Mann-Whitney test. **C**. PAC2 cells were transfected with memCherry and stained with an antibody against Ror2. **D**. PAC2 cells were transfected with indicated constructs and stained with indicated antibodies. High-magnification images indicate co-localization of Wnt5a/b and Ror2 on cytonemes (yellow arrows). **E**. Shows live imaging of co-transfected PAC2 cells with indicated markers at 24 h of post-transfection. Wnt5b, Ror2, and Wnt5b/Ror2 clusters can be observed in the non-transfected neighboring cells (yellow arrow). R, receiving cells; P, producing cells; scale bar 5µm. **F, G**. Wild-type PAC2 were co-cultivated with ROR2^CRISPR^ cells for 24h, fixed, and stained with an antibody against Ror2 and Phalloidin-405. Scale bar 10µm. **G** shows high magnification images of the yellow box indicated in **F**. Orange arrows indicate Ror2 puncta at the plasma membrane of WT PAC2 cells. Yellow arrows indicate Ror2 puncta in the adjacent Ror2 ^-/-^PAC2 cells. Blue arrows indicate filopodia protrusions. **H**. Time series of cytoneme-based transportation of Wnt5b/Ror2 in wild-type zebrafish embryos from 0’00“to 3’50”. Embryos were injected with mRNA for Wnt5b-GFP, Ror2-mCherry, and mem-BFP at the 8-cell stage in one blastomere to generate local clones and imaged live at 6 hpf. R, receiving cells; P, producing cells; scale bar 10 µm. Blue arrows indicate Wnt5b/Ror2 clusters. Yellow arrows indicate cytonemes. **I, J**. PAC2 zebrafish fibroblasts transfected with indicated constructs and imaged live at 24 h of post-transfection. Yellow arrows co-localization of indicated markers with Wnt5b or Ror2 on cytonemes. Scale bar 5µm.

We then investigated this co-transport of Wnt5b together with Ror2 in zebrafish embryos (Fig.1 H; Suppl. Fig. 1 E, F; Suppl. Video 1). In less than 4min, we observed the formation of a cytoneme, the hand-over of Wnt5b-GFP and Ror2-mCherry, and the subsequent retraction of the protrusion. The transport of Wnt5b-GFP/Ror2-mCherry was accompanied by the independent membrane marker mem-BFP, suggesting a transfer of the entire cytoneme tip vesicle to the receiving cell. Finally, we further characterized these Wnt5b/Ror2 positive protrusions and found that they also carry the filopodia tip markers, the unconventional MyoX, and N-WASP (Fig. 1I, J). Therefore, we conclude that Wnt5b and Ror2 can be loaded on signaling filopodia, referred to as Wnt/PCP cytonemes hereafter. After elongation and contact formation of the Wnt/PCP cytoneme, the Wnt5b/Ror2 positive cytoneme tip is delivered to the neighboring cell during zebrafish gastrulation.

### Wnt5b/Ror2 forms a tight complex during transport

Based on the surprising finding that Wnt5b/Ror2 can be handed over together to neighboring cells, we aimed to characterize the intermolecular interaction within this ligand-receptor complex in the living zebrafish embryo. Therefore, we adapted two *in vivo* techniques: Fluorescence Lifetime Imaging Microscopy - Förster Resonance Energy Transfer (FLIM-FRET) and Fluorescence Cross-Correlation Spectroscopy (FCCS). First, we wanted to describe the complex with a high spatial and temporal resolution regarding the differentially tagged fluorescent components. Therefore, we chose FLIM-FRET because it is independent of the fluorophore concentration, the excitation efficiency, and the impact of light scattering - a prerequisite for analysis *in vivo* (Suppl. Fig. 2A, B). First, we measured the FRET efficiency of two GPI-anchored, lipid-raft associated fluorescent proteins, memGFP, and mem-mCh, as a positive control and compared these measurements to our negative control, a memGFP a cytosolic mCherry (cyto-mCh; Fig. 2A, B). We found that memGFP/mem-mCh and memGFP/cyto-mCh show significant differences concerning the GFP lifetime (2.24ns versus 2.41ns) and the FRET efficiency (11.1% versus 4.6%), and the distance between the fluorophores (74.7Å versus 92.7Å), respectively (Fig. 2G-I). Next, we measured the lifetime, the FRET efficiency, and the distance of the donor Wnt5b-GFP and the N-terminally tagged acceptor molecule mCherry-Ror2, which is biological active (Suppl. Fig. 4C). To this end, we visually selected appropriate Wnt5b-GFP clusters in cells of the producing cell, on cellular protrusions emitting from these source cells, and in the adjacent receiver cell (Fig. 2C-E, blue arrows). At all three locations, we measured a similar GFP lifetime of ca. 2.16-2.19ns, FRET efficiency of ca. 11.3-13%, and distance of 75.7-75.0Å between Wnt5b-GFP and mCherry-Ror2 in clusters in the producing cell, on cytonemes, and in the receiving cells (Fig. 2C-E, G-I). Finally, we measured the GFP lifetime, FRET efficiency, and the distance between Wnt5b-GFP and a Ror2, which carries a C-terminal mCherry tag. In this setting, the increased space between the fluorophores and the membrane between the fluorophores acting as an insulator should reduce FRET efficiency significantly. (Suppl. Fig. 2D). Indeed, we measured a GFP lifetime of 2.40ns, FRET efficiency of only 3.6%, and distance between the fluorophores of 101.8Å (Fig. 2F, G-I), comparable to the memGFP/cyto-mCh control. This data set indicates that mCherry-Ror2 binds to Wnt5b-GFP and that this complex maintains its structural assembly concerning the packaging of the fluorescently tagged constructs during transport.

**Figure 2.**
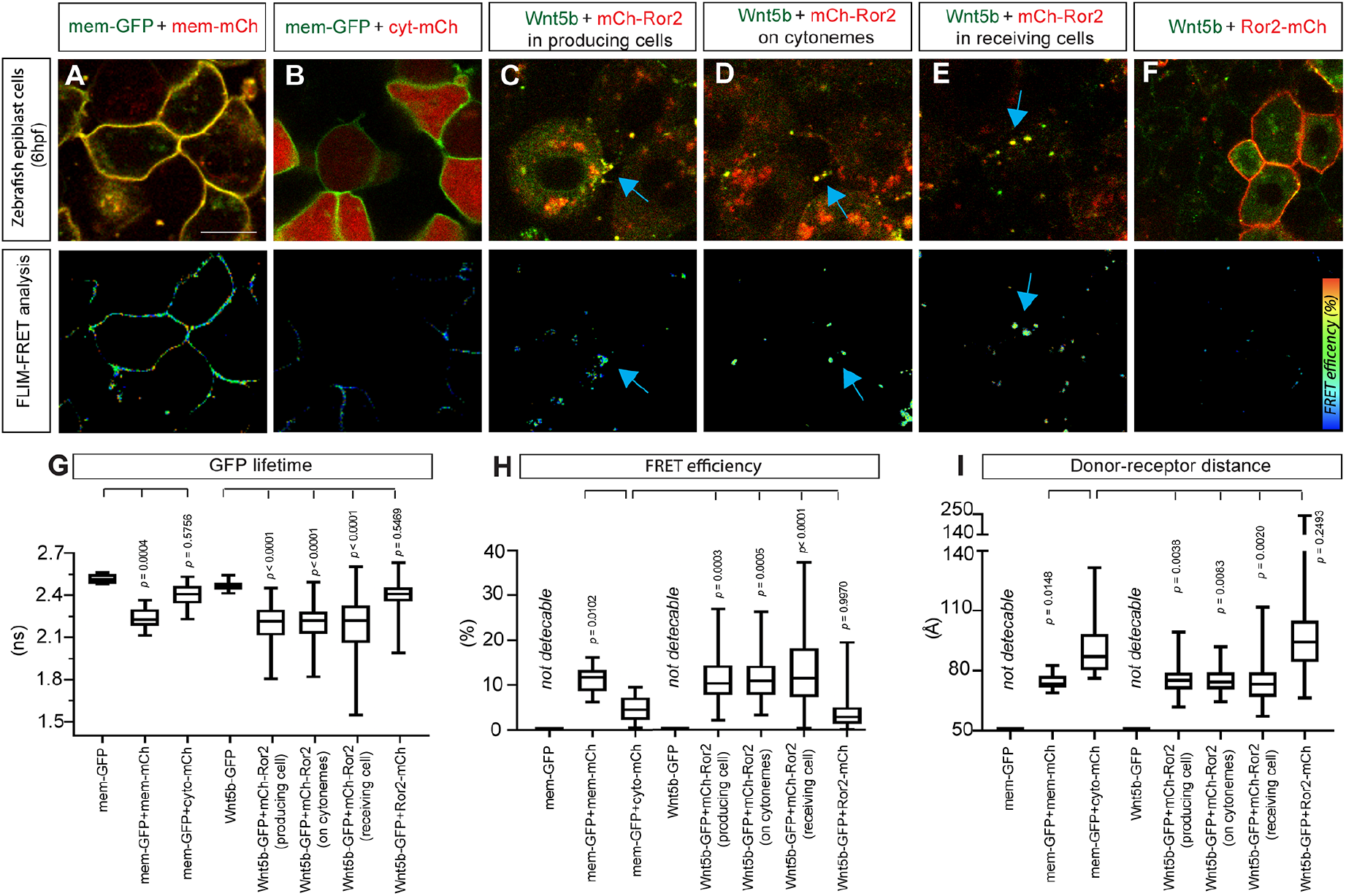
*In vivo* FLIM-FRET imaging reveals maintenance of Wnt5b/Ror2 complex cohesiveness during transport. **A-F**. Wild-type zebrafish embryos were injected with the mRNA for the indicated constructs at the 8-cell stage in one blastomere to generate local clones and imaged live at 6 hpf. The representative fluorescence images and corresponding FLIM-FRET analysis are shown. Scale bar 10µm. The blue arrows indicate the Wnt5b/Ror2 clusters with a high FRET efficiency. **G-I**. Quantification of fluorescent GFP lifetime (n=8, 17, 25, 21, 36, 23, 33, 49, n= numbers of cells), FRET efficiency and donor-receptor distance (n=0, 17, 24, 0, 36, 22, 30, 43, n= regions of interests). Significance is calculated by a one-way ANOVA test plus Tukey’s multiple comparisons test. *p*-values shown on the graph are compared to the negative control (memGFP + cyto-mCh).

### The binding affinity between Wnt5b and Ror2 is maintained during transport

FCCS can determine the affinity, quantified by the equilibrium dissociation coefficient, K_D_, as a key parameter for the specificity of ligands and receptors in a signaling pathway. We thus performed a set of FCCS experiments to measure molecular interactions of fluorescently tagged signaling components in the Wnt5b/Ror2 complex in the living zebrafish embryo (Suppl. Fig. 3A-D). Based on the calibration measurements of the *in vivo* FCCS measuring system, we can perform fluctuation recordings in a predefined volume of 0.65*10^−9^ nm^3^ with a recording time of 10sec (Suppl. Fig. 3E. F). First, we determined the cross-correlation of mem-GFP and an αGFP-nanobody coupled to mCherry (Vhh-mCherry) at the plasma membrane of mesenchymal cells of living zebrafish embryos. Here, we find a strong cross-correlation between memGFP and Vhh-mCherry. For this positive control, we measured a dissociation constant K_D_ of 229nM (Fig. 3A, H). Then, we determined the binding affinity of a memGFP to a cyto-mCh, as a negative control. In fact, we did not observe cross-correlation and calculated a K_D_ value of over 9301nM. Based on these values, we measured cross-correlation for Wnt5b-GFP and Ror2-mCherry *in vivo*. As our FCCS setup allows fast measurements in tiny volumes, we could also determine the cross-correlation at various subcellular localisations, here the cell membrane of the Wnt5b-GFP/Ror2-mCherry producing cells, on cytoneme tips, and in clusters in the receiving cells in the living zebrafish embryo. We determined the K_D_ values for Wnt5b/Ror2 in the producing cell, on cytonemes, and in the receiving cell from 311-476nM (Fig. 3H). Similar to the FLIM-FRET analysis, we found an equal distribution of the individual K_D_ measurements at the different subcellular localisations, suggesting that the ligand-receptor complex is maintained during translocation from the producing cell to the receiving cell (Fig. 3C-E, H). As a negative control experiment, we measured the binding affinity of Wnt5b to Ror2 lacking the Wnt binding domain (CRD domain; Oishi et al., 2003). Similar to our memGFP/cyto-mCh negative control, we did not observe cross-correlation, and we calculated a K_D_ of over 6678nM, suggesting a very low binding affinity (Fig.3F, H). Based on the *in vivo* FLIM-FRET and FCCS analysis, we assume that the Wnt5b/Ror2 complex is maintained during transport. To challenge the integrity of the complexes, we over-expressed Wnt5b-GFP and untagged Ror2 together in the source cell and overexpressed Ror2-mCherry in the receiving cell. As FCCS is very sensitive, an exchange of even a few Ror2-mCherry molecules with untagged Ror2 molecules could be detected. However, in this experiment, we find no cross-correlation between Wnt5b-GFP and Ror2-mCherry and calculated a K_D_ from 3560nM, suggesting that the Wnt5b-GFP/Ror2 complex stays largely intact, even if there is an excess of unbound Ror2 in the target cell.

**Figure 3.**
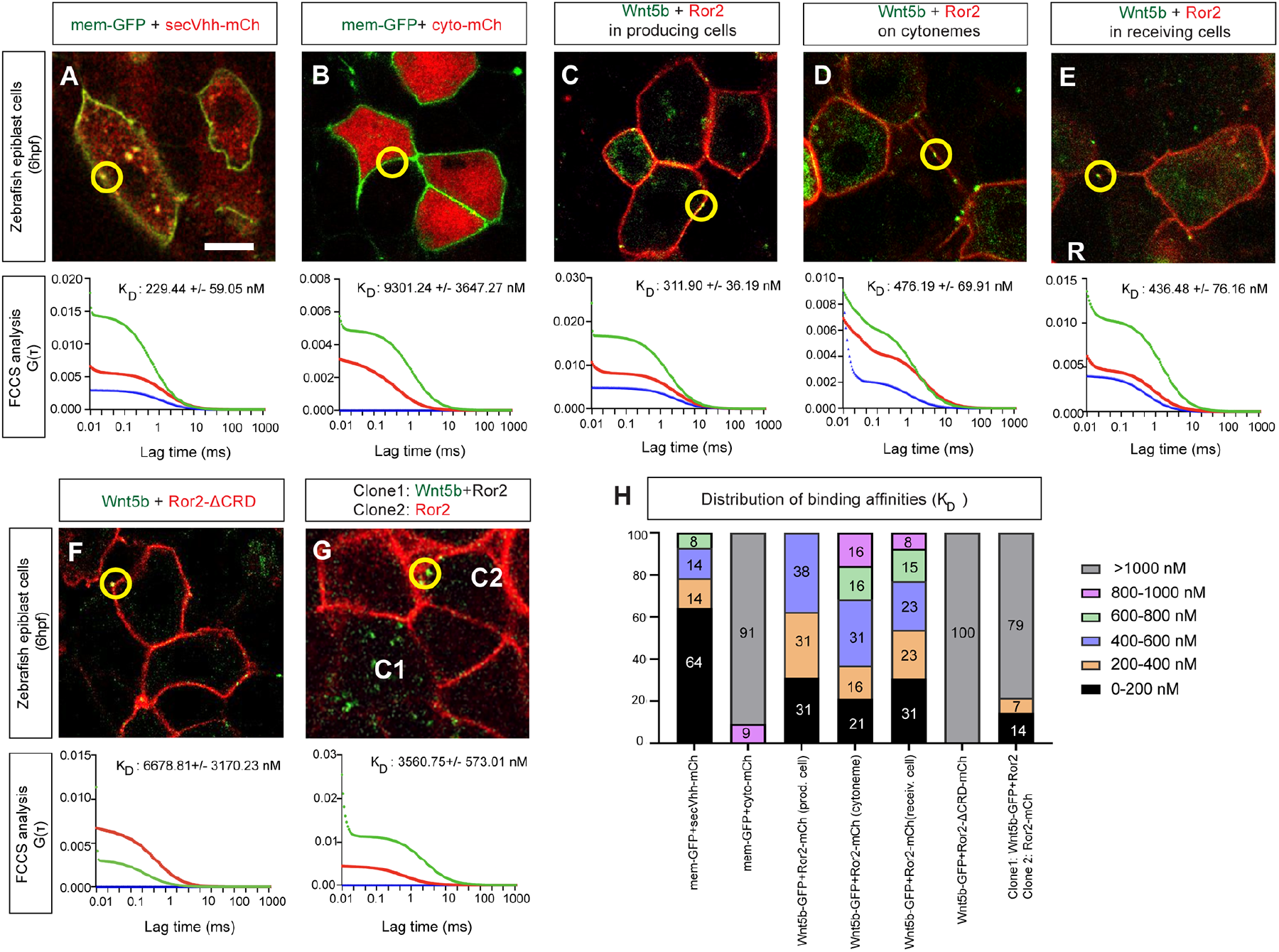
*In vivo* fluorescence cross-correlation spectroscopy (FCCS) analysis suggests that binding affinities of Wnt5b to Ror2 are maintained during complex transport. **A-F**. Wild-type zebrafish embryos were injected with the mRNA for the indicated constructs at the 8-cell stage in one blastomere to generate local clones and imaged live at 6 hpf. The representative fluorescence images and corresponding FCCS analysis were displayed. P, producing cells; R, receiving cells. Scale bar 10µm. **G**. Wild-type zebrafish embryos were injected with the mRNA for Wnt5b-GFP and Ror2 in one blastomere and Ror2-mCherry mRNA in the neighboring blastomere at the 8-cell stage to generate local clones and imaged live at 6 hpf. C1 indicates the Wnt5b-GFP/Ror2 positive clone, and C2 indicates a clone expressing Ror2-mCherry. Yellow circles indicate the measuring spot during FCCS. The auto-correlation curves for Wnt5b-GFP and Ror2-mCherry are shown in red and green, respectively. The cross-correlation curve is shown in blue. The corresponding average dissociation constant (K_D_) is indicated on the graph. **H**. Breakdown of the percentage of binding affinities into 0-400, 400-600, 600-800, 800-1000, >1000 nM categories (n= 15, 11, 17, 19, 14, 8, 17, n= numbers of filtered measurements). The individual percentage is shown on the bar.

### Wnt5b/Ror2 complexes are active in the source cells and on cytonemes

Although we observed the transfer of Wnt5b/Ror2 clusters to the receiving cells, we do not know whether these clusters can signal during assembly, transport, and at their final destination. First, we asked which processes require Wnt5b/Ror2 activity in the source cells and, more specifically, whether Wnt5b/Ror2 affects cytoneme formation per se (Fig. 4). To tackle this question, we overexpressed the indicated PCP constructs in clones in the zebrafish embryo and quantified the length and the number of filopodia per cell. We found that activation of Wnt5b, Ror2, and Wnt5b/Ror2 led to the formation of fewer but much longer filopodia (Fig. 4A-D, I, J). IRSp53, a multidomain BAR protein, and its binding partner, the small GTPase Cdc42, promote filopodia formation (Kast et al., 2014). We asked if the Wnt/PCP-induced filopodia are dependent on these filopodia regulators. To block cytoneme formation in zebrafish without interfering with Wnt signaling, we used the mutated protein IRSp53^4K^ and the dominant-negative Cdc42^T17N^ (Brunt et al., 2021; Stanganello et al., 2015). We found that the formation of the Wnt/PCP-induced long cytonemes can be significantly reduced by co-expression of IRSp53^4K^ or Cdc42^T17N^ (Fig. 4E-H, I). In parallel, we investigated the distribution of filopodia numbers per cell (Fig. 4J). We observed that activation of Wnt/PCP led to a downregulation of the number of filopodia. We observed a similar phenotype when we blocked filopodia formation by expression of IRSp53^4K^ and Cdc42^T17N^. Therefore, we conclude that Wnt/PCP is required to induce long cytonemes in the source cell.

**Figure 4.**
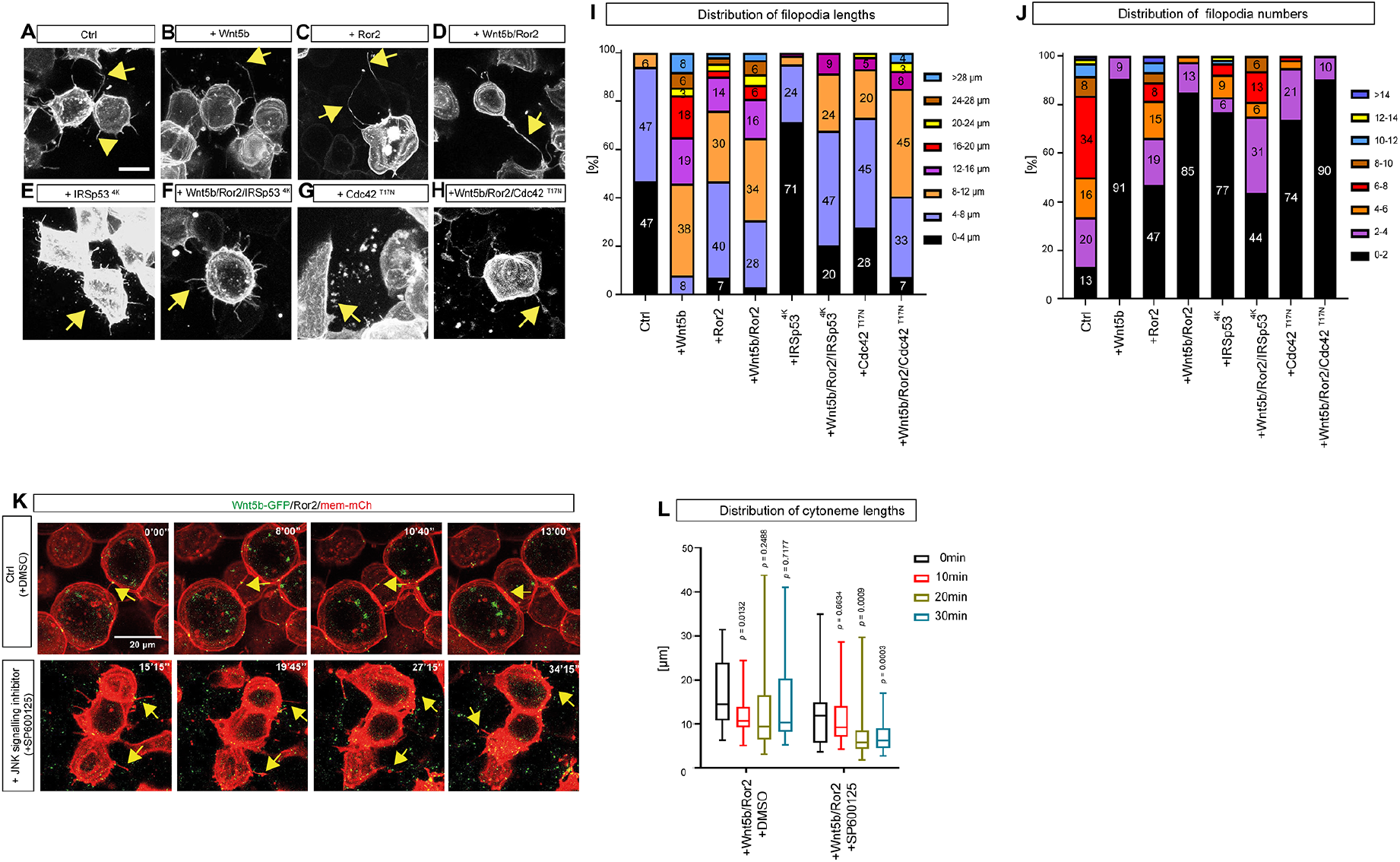
Wnt5b/Ror2/JNK signaling promotes the formation of long cytonemes. **A-H**. Confocal images of wild-type zebrafish embryos injected with indicated constructs at 6hpf. Yellow arrows indicate filopodia. **I**. The quantitative analysis of filopodia length in zebrafish embryos, breaking down the percentage into 0-4, 4-8, 8-12, 12-16, 16,20, 20-24, 24-28, >28 μm categories. (n=367, 63, 113, 68, 41, 59, 119, 27, n= numbers of measurements). Scale bar 10µm. **J**. The quantitative analysis of filopodia numbers per cell in zebrafish embryos, breaking down the percentage of numbers per cell into 0-2, 2-4, 4-8, 8-10, 10-12, 12-14, >14 categories. The individual percentage is shown on the bar. **K**. Time series of cytoneme in wild-type zebrafish embryos. Embryos were injected with Wnt5b-GFP/Ror2/mem-mCherry in the clone. At 6hpf, the embryos were treated with DMSO or JNK inhibitor SP600125 from 0‘00“to 30’00” and imaged live. Yellow arrows indicate the retraction of cytonemes at the corresponding time points. **L**. The quantitative analysis of cytoneme length at the different time points in zebrafish embryos was treated as indicated in (**K**). (Wnt5b/Ror2: n=25, 23, 16, 13; Wnt5b/Ror2/SP600125: n= 35, 34, 38, 54, n= numbers of cytonemes).

Filopodia formation is dependent on JNK signaling (Bosch et al., 2005; Martin-Blanco et al., 2000) and, specifically, the upkeep of Wnt8a cytonemes (Brunt et al., 2021). Therefore, we asked if the extension of Wnt/PCP-cytoneme depends on continuous JNK signaling. We found the long Wnt5b/Ror2-positive cytonemes fragment minutes after treatment with the JNK signaling inhibitor SP600125 (Fig.4K, L; Suppl. Video 2). These experiments suggest that Wnt/PCP signaling is required twofold: Firstly, Wnt/PCP signaling induces Wnt5b/Ror2-bearing cytonemes. Secondly, we further observe that continuous Wnt5b/Ror2/JNK signaling is crucial for shaping long cytonemes.

### Transferred Wnt5b/Ror2 complexes activate paracrine JNK signaling

Next, we asked whether the transferred Wnt5b/Ror2 complexes maintain their activity in the target cells. Wnt5b/Ror2-mediated PCP signaling activates JNK signaling in the zebrafish gastrula (Brunt et al., 2021). Therefore, we used the JNK kinase translocation reporter, JNK KTR-mCherry, to monitor alteration in the JNK signaling cascade (Fig. 5). JNK-KTR-mCherry localizes to the nucleus (N) in its dephosphorylated state (low JNK activity, Suppl. Figure 4A). Upon activation by phosphorylation, it shuttles to the cytoplasm within minutes (C, high JNK activity). The C: N ratio can then be calculated as an indicator of JNK signaling strength in near real-time. To address the function of paracrine Wnt/PCP signaling in zebrafish embryos, we ubiquitously overexpressed the reporter of JNK signaling. Within this tissue, we generated small clones, which expressed signaling components and cytoneme regulators and measured JNK signaling strength in the neighboring cell rows (Suppl. Fig. 4B). We observed that a control clone expressing memGFP does not interfere with JNK signal activation. Instead, the adjacent cells display low basal levels of JNK activity (Fig. 5A). Next, we observed that a Wnt5b-positive clone activates JNK signaling; however, predominantly in the cells adjacent to the clone (Fig. 5B, arrow). Furthermore, clonal Ror2 expression strongly activates the JNK reporter concentration-dependent manner. This is also influenced by distance from the clone (Fig. 5C, E). Finally, we show that clonal Wnt5b/Ror2 expression activates JNK signaling, even on a broader signaling range (Fig. 5D, F).

**Figure 5.**
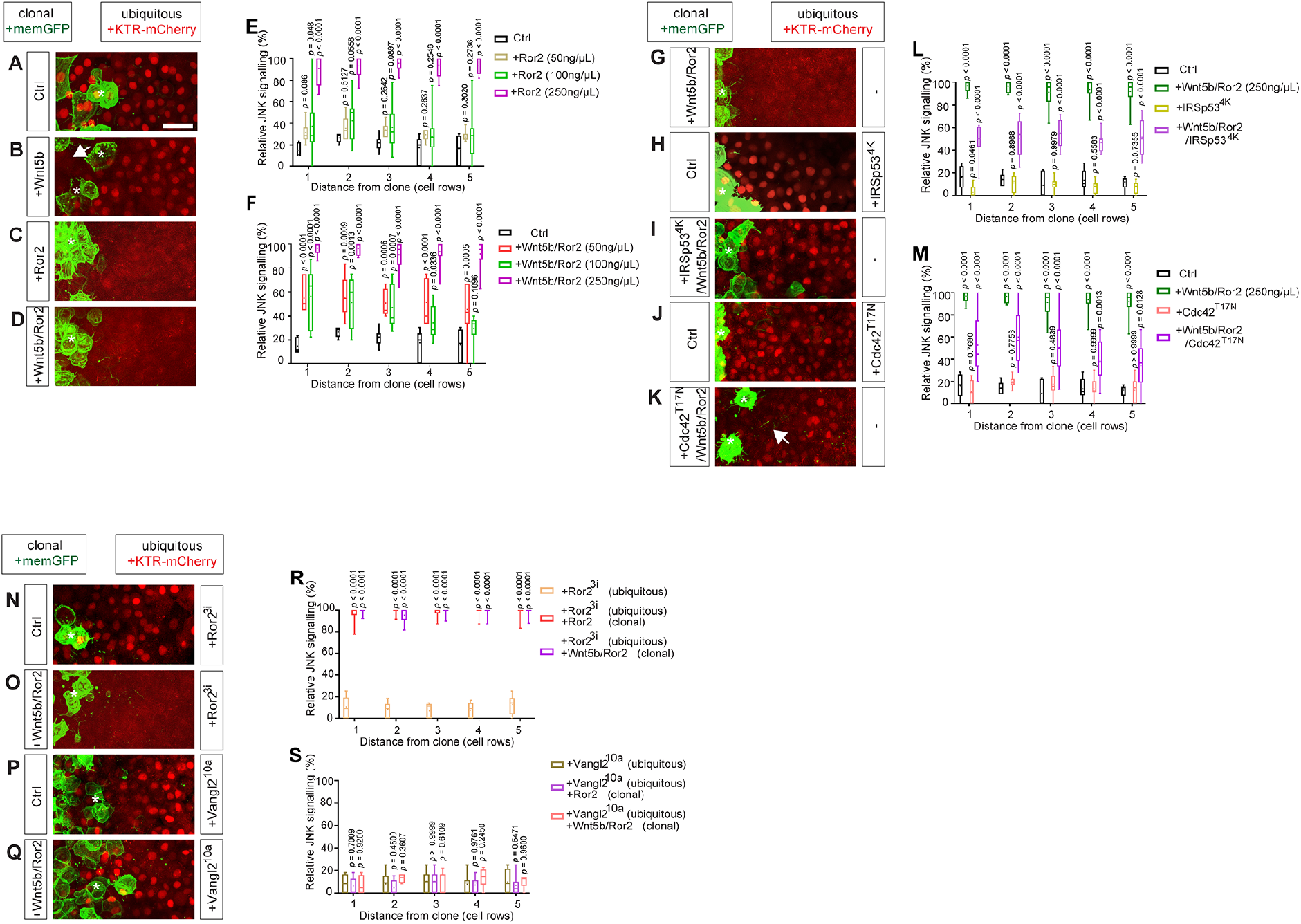
Wnt5b/Ror2 expressing clones activate autocrine and paracrine JNK signaling. **A-D, G-K, N-Q**. Wild-type zebrafish embryos were ubiquitously injected with KTR-mCherry and indicated constructs (ubiquitous). At the 8-cell stage, one blastomere was injected with the indicated constructs (clonal) to generate clones at the embryonic margin and imaged live at 6 hpf. White stars indicate the signaling producing clonal cells expressing memGFP. Scale Bar 20µm. **E-F**. Relative JNK signalling within 5 cell rows distance from clone when injected with different amount of Ror2 (n= 7, 7, 11, 10, n= numbers per embryo) and Wnt5b/Ror2 (n= 7, 7, 12, 12, n= numbers per embryo). **L, M**. Relative JNK signalling within 5 cell rows distance from clone when injected with IRSp53^4k^ (n= 6, 12, 8, 9, n= numbers per embryo) and Cdc42^T17N^ (n= 6, 12, 11, 14, n= numbers per embryo). **R-S**. Relative JNK signalling within 5 cell rows distance from clone when injected with Ror2^3i^ (n= 9, 10, 11, n= numbers per embryo) and Vangl2^10a^ (n= 11, 11, 6, n= numbers per embryo). Significance is calculated by two-way ANOVA together with Dunnett’s multiple comparisons test. *p*-values as indicated.

Next, we asked if paracrine Wnt5b/Ror2 signaling requires cytonemes. To block cytoneme formation, we used the dominant-negative mutants IRSp53^4K^ and Cdc42^T17N^. We found that ubiquitous overexpression of IRSp53^4K^ does not interfere with JNK signaling (Fig. 5H). Next, we blocked cytoneme formation exclusively in the clonal cells by overexpressing IRSp53^4K^ together with Wnt5b/Ror2 in the clones (Fig. 4I). We observed that IRSp53^4K^/Wnt5b/Ror2 positive clones significantly reduced the capability to activate paracrine PCP/JNK signaling compared to Wnt5b/Ror2 clones (Fig. 5G, L). To validate our findings, we blocked actin polymerization by overexpressing the Cdc42^T17N^ (Nalbant et al., 2004), a prerequisite for Wnt cytoneme formation in zebrafish (Stanganello et al., 2015). Similar to the ubiquitous expression of IRSp53^4K^, Cdc42^T17N^ overexpression did not interfere with JNK signaling (Fig. 5J). However, Wnt5b/Ror2 clones, which co-express Cdc42^T17N^, had a reduced ability to activate paracrine JNK signaling (Fig. 5K, M). Notably, cells in the close vicinity of the clone still showed low JNK activation (white arrow), suggesting that Cdc42^T17N^ is less potent in blocking cytoneme formation than IRSp53^4K^. In summary, these data indicate that paracrine Wnt/PCP signaling is strongly dependent on cytoneme appearance in zebrafish development.

Next, we asked if the transferred Wnt5b/Ror2 complex remains active in the target cells. Therefore, we blocked Wnt/PCP signaling in the receiving cell by overexpressing a dominant-negative form of the Ror2 receptor, the kinase-dead mutant Ror2^3i^ (Hikasa et al., 2002), and the dominant-negative form of Vangl2, the all-phospho Vangl2 mutant S5∼17A∷ S76∼84A, in which 10 Ser/Thr are replaced by Ala (Vangl2^10A^) (Gao et al., 2011). We found that overexpression of Ror2^3i^ blocks JNK signal activation (Fig. 5N). However, this blockage of Ror2/PCP signaling by ubiquitous overexpression of Ror2^3i^ can be overcome by paracrine Wnt/PCP from the clone (Fig. 5N, O, R). We found that even an excess of kinase-dead receptors cannot hinder signal transduction of the transferred Wnt5b/Ror2 complexes, suggesting that the transmitted Wnt5b/Ror2 complexes remain assembled and active in the receiving cell, consistent with our *in vivo* FLIM-FRET and FCCS analysis. Wnt5b/Ror2 signaling requires Vangl2 to activate JNK signaling (Brunt et al., 2021; Gao et al., 2011). Therefore, we overexpressed the dominant-negative mutant Vangl2^10A^ (Brunt et al., 2021; Gao et al., 2011) in the target cells, blocking Wnt/PCP signaling. We found that Vangl2 disturbs Wnt/PCP mediated JNK signaling, even in cells in close vicinity of the Wnt5b/Ror2 clone (Fig. 5P, Q, S). This data suggests that active Wnt5b/Ror2 are handed over to the receiving cells to activate JNK signaling within these cells. The transferred Wnt5b/Ror2 complex requires Vangl2 for transducing Wnt/PCP signaling in the receiving cell.

### Cytonemal-delivered Wnt5b/Ror2 complexes can block Wnt/β-catenin signaling

We analyzed the paracrine signaling function of Wnt/PCP clones with regard to endogenous gene expression in zebrafish gastrulation. Besides activation of JNK signaling, Wnt/PCP signaling acts antagonistically to the Wnt/β-catenin pathways, and it has been shown that Wnt/PCP signaling effectively represses Wnt/β-catenin target gene expression (Niehrs, 2012; van Amerongen and Nusse, 2009). Therefore, we generated small clones marked with a lineage tracer in the zebrafish embryonic margin and subjected these embryos to *in-situ* hybridization with a probe against the bona fide Wnt/β-catenin target gene *lef1*. At 6hpf, *lef1* is expressed predominantly in the embryonic margin (Fig. 6A). On the other hand, the expression of Wnt5b, Ror2, or Wnt5b/Ror2 reduces the expression of *lef1* within the clonal cells (Fig. 6B-D, E) significantly. In addition, Ror2 and Wnt5b/Ror2 also strongly reduce the expression in the embryonic margin, whereas Wnt5b expression reduces *lef1* expression in a halo around the clone (Fig. 6B-D, F).

**Figure 6.**
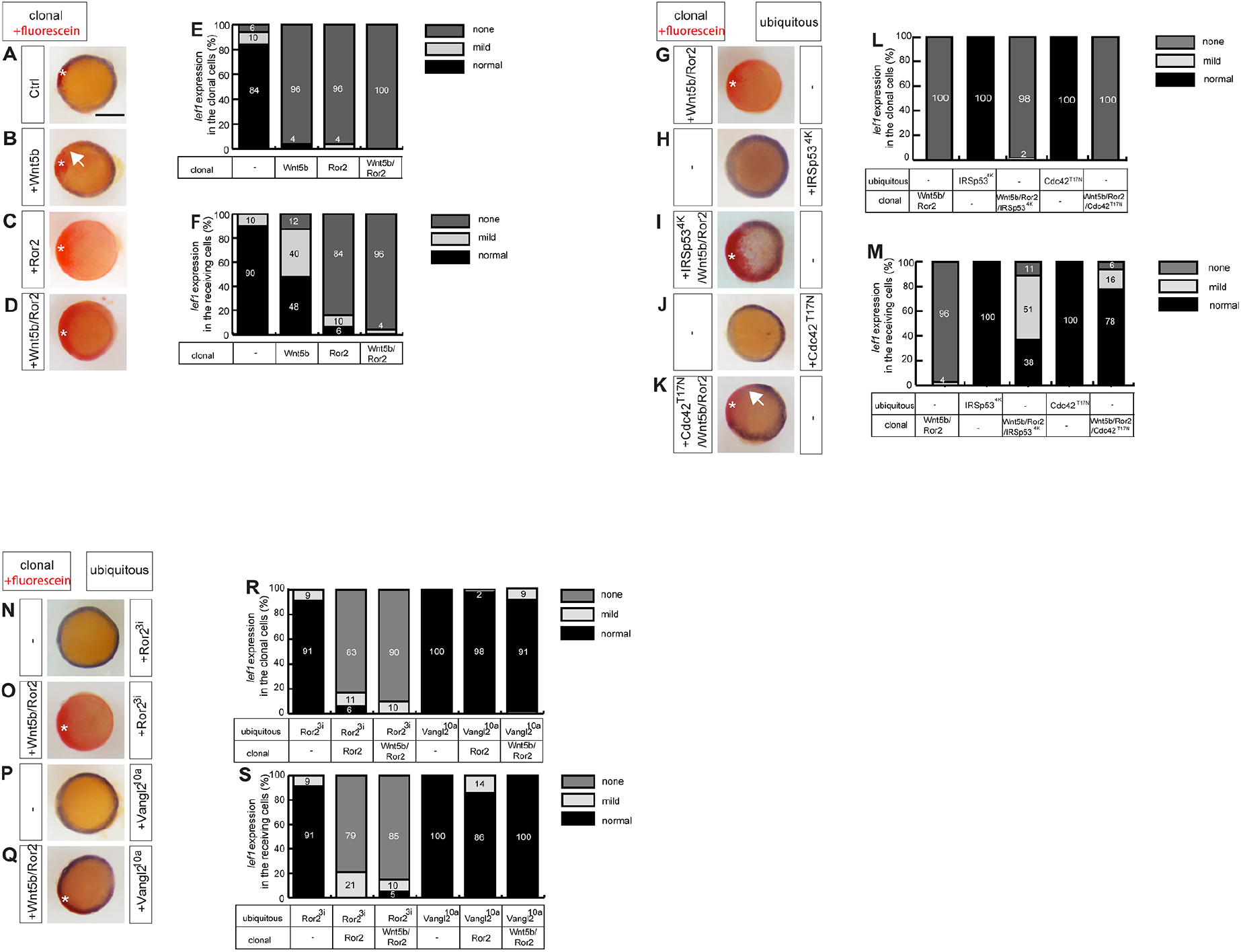
Wnt5b/Ror2 complexes repress the expression of the Wnt/β-catenin target gene *lef1*. **A-D, G-K, N-Q**. Embryos injected with the mRNA for the indicated constructs at the one-cell stage (ubiquitous) or at the 8-cell stage in one blastomere to generate local clones (clonal) were subjected to an *in situ* hybridization analysis against *lef1* at 6hpf. The embryos were stained and mounted with the animal pole up. Purple staining around the margin indicates the expression of *lef1*, and red staining (white star) indicates the site of the injected clonal cells. Clonal cells were co-injections with Mini-Emerald as a lineage tracer. Scale bar 200µm. **E-F, L-M, R-S**. Classification of *lef1* expression in the clonal cells and the receiving cells. The expression is classified according to the severity of the downregulation of the *lef1* expression into none (e.g., C), mild (e.g., B), and normal (e.g., A). Numbers in bars represent the percentage of total embryos in each group.

Next, we asked if cytoneme-based transport is required for the observed repression of Wnt/β-catenin expression. Therefore, we overexpressed Wnt5b/Ror2 in the clone and observed a strong repression of *lef1* expression in the clone as well as in the embryonic margin (Fig. 6G, L). Next, we blocked cytoneme formation in these Wnt5b/Ror2 clones by co-expression of IRSp53^4K^ or Cdc43^T17N^. Ubiquitous expression of IRSp53^4K^ or Cdc43^T17N^ indicated that the function of these cytoneme inhibitors is independent of the induction of Wnt/β-catenin signaling, shown by the expression of the *lef1* in the embryonic margin (Fig. 6H, J). After blocking cytoneme formation in Wnt5b/Ror2 expressing clones by IRSp53^4K^ or Cdc43^T17N^, we found that *lef1* expression can be observed in the embryonic margin, whereas *lef1* expression is inhibited within the clone (Fig. 6I, K), suggesting that cytonemes are required for paracrine Wnt/PCP signal dissemination. In contrast, cytonemes are dispensable for autocrine signal activation.

Finally, we addressed whether signaling of the transferred Wnt5b/Ror2 complexes requires endogenous signaling components. Therefore, we blocked Wnt/PCP signaling by ubiquitous expression of Ror2^3i^ and Vangl2^10a^, which did not interfere with *lef1* expression in the embryonic margin (Fig. 6N, P). Wnt5b/Ror2-expressing clones can overcome the Wnt/PCP block of Ror2^3i^, suggesting that Wnt5b/Ror2 remain active and repress Wnt/β-catenin signaling in the receiving cells (Fig. 6O, R, S). However, blockage of Wnt/PCP signaling by the dominant-negative Vangl2 mutant Vangl2^10a^ blocks repression of *lef1* expression in the clone and the margin (Fig. 6Q, R, S), suggesting that Vangl2 is an essential mediator of Wnt/PCP signaling for endogenous Wnt/Ror2 complexes as well as for the transferred complexes in the zebrafish embryo.

### Paracrine signaling of Wnt5b/Ror2 regulates C&E during zebrafish embryogenesis

Next, we addressed whether convergence and extension (C&E), a hallmark of active Wnt/PCP signaling, is affected by cytoneme-mediated dissemination of active Wnt/PCP complexes. C&E are crucial for the migration of the axial mesoderm towards the midline in the developing zebrafish embryo. Therefore, we used the transgenic zebrafish line, Gsc-GFP^CAAX^, which expresses a membrane-tethered form of GFP under the control of the Gsc promoter in the notochord (Smutny et al., 2017). Similar to the previous experiments, we generated small clones in the lateral mesoderm expressing the indicated PCP constructs (Suppl. Fig. 4D). At 10hpf, we mounted the embryos and performed confocal microscopic imaging of the developing notochord to quantify the width of the notochord. We found that clones expressing Ror2 (Fig. 7B, G) or clones expressing Wnt5b and Ror2 (Fig. 7D) led to a significant broadening of the notochord anlage in comparison to a mCherry expressing ctrl cone (Fig. 7A). To address whether these effects are based on Wnt/PCP cytonemes emitting from the clonal cells, we reduced the capability of the clonal cells to form cytonemes. We found that these Wnt/PCP induced defects can be partially rescued by co-expressing IRSp53^4K^ and thus inhibiting cytoneme formation (Fig. 7C, E). However, IRSp53^4K^ expressing clones cause hardly any notochord alterations (Fig. 7F). The increased width of the notochord could be explained by a less tightly packed tissue due to less elongated and more rounder NC cells. Therefore, we further determined the cell circularity as the ratio between the cell diameter of the Gsc-GFP positive cells parallel to the body axis versus the cell diameter perpendicular to the body axis Suppl. Figure 4E). Indeed, we find that notochord cells display a higher circularity if they are in close vicinity of a Wnt5b/Ror2 (Fig. 7H). Again, these Wnt/PCP induced defects can be partially rescued by co-expressing IRSp53^4K^ and thus inhibiting cytoneme formation.

**Figure 7.**
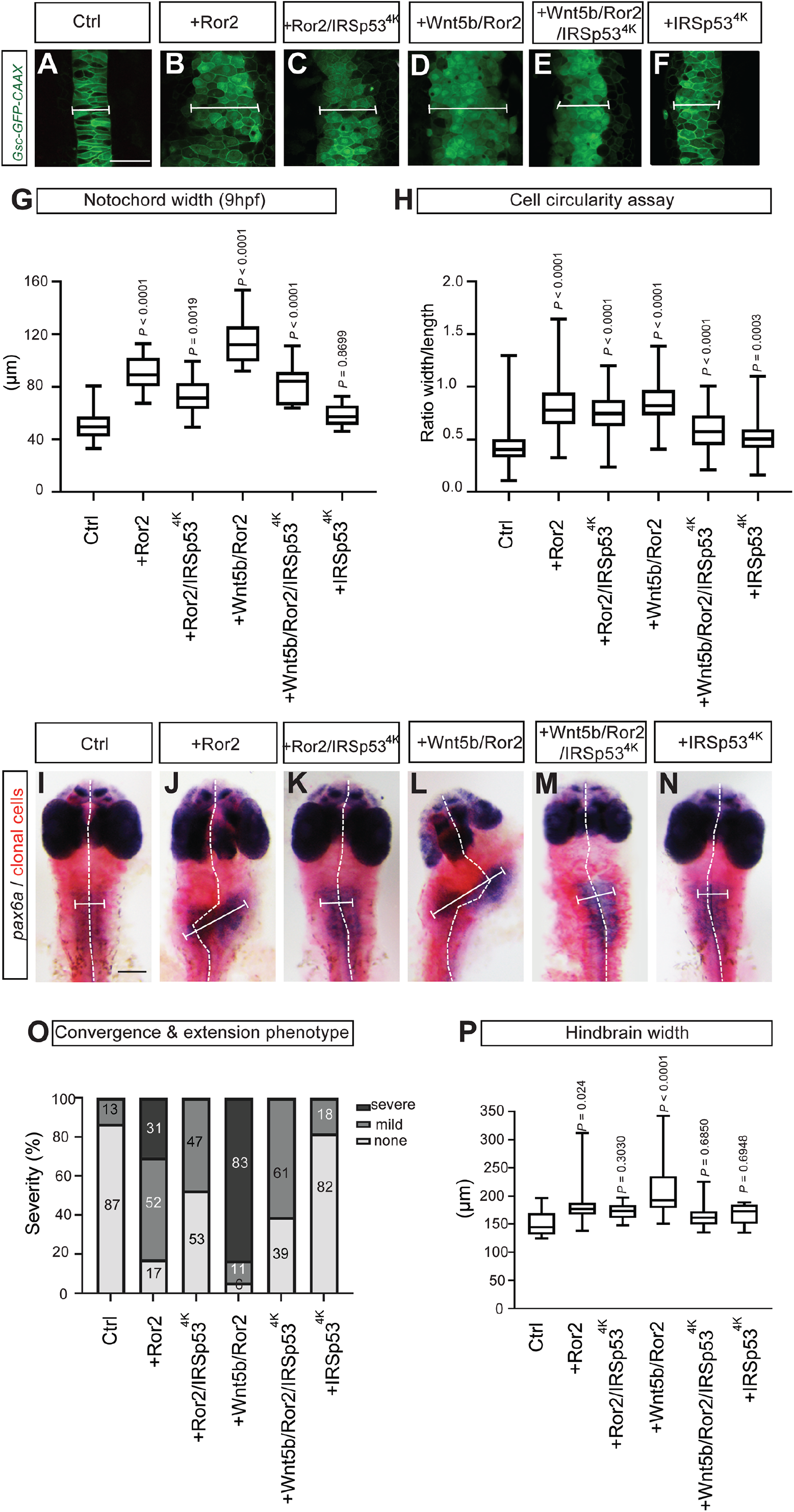
Alterations of cytoneme-mediated dissemination of Wnt5b/Ror2 complexes affect convergence and extension in the zebrafish embryo. **A-F**. *Tg (−6gsc: EGFP –CAAX)* zebrafish embryos were injected with the mRNA for the indicated constructs at the 8-cell stage in one blastomere to generate local clones imaged live at 10hpf. Scale bar 50µm. The white line indicates the notochord width. **G**. The notochord width was measured in embryos (n= 20, 13, 12, 8, 12, 9, n= numbers per embryo). **H**. Cell circularity was measured within each embryo (n= 173, 292, 194, 200, 149, 176, n= numbers per cell). A perfect circular cell has a circularity of 1.0, while below or above 1.0 indicates a noncircular, elongated shape. **I-N**. Wild-type zebrafish embryos with the mRNA for the indicated constructs plus Mini-Emerald as lineage tracer at the 8-cell stage in one blastomere to generate local clones and fixed at 30 hpf for *in situ* hybridization against a probe for *pax6a*. White dashed lines indicate the course of the midline, and the white bars indicate the width of the hindbrain domain. Scale bar 200µm. **O**. The quantitative analysis of phenotype severity in zebrafish embryos. The phenotype severity is classified into none (e.g., I), mild (e.g., J), and severe (e.g., L). I, J, and L are representative examples of the different severity classifications, respectively. Numbers in bars represent the percentage of total embryos in each group. **P**. Hindbrain width was measured in embryos. N= 13, 23, 19, 18, 23, 11, n= numbers per embryo. Significance is calculated by a one-way ANOVA test plus Tukey’s multiple comparisons test.

At 32hpf, we subjected these embryos to an *in situ hybridization* against *pax6a* to analyze the effects of the Wnt/PCP clones in the neuroectoderm. Similar to the notochord analysis, we found that Ror2 expressing clones, and even stronger Wnt5b/Ror2 expressing clones, interfere with C&E by increasing the width of the neuroectoderm in the developing embryo (Fig. 7I, J, L, O, P). However, again co-expression of IRSp53^4K^ partially rescues the effect of the Wnt/PCP clones (Fig. 7K, M), and IRSp53^4K^ clones have no detectable influence on the narrowing of the neuroectoderm (Fig. 7N). Based on these data sets, we conclude that paracrine Wnt5b/Ror2 complexes - disseminated by cytonemes - play a pivotal role in regulating local C&E in zebrafish development.

## Discussion

Wnt/PCP signaling orchestrates essential processes in vertebrate development, such as cell polarity and tissue migration. To this end, Wnt proteins can act as autocrine and paracrine signaling proteins (Routledge and Scholpp, 2019). For example, Wnt5b ligands can form gradients to determine the spatial identity and influence the behavior of the target cells in the mouse limb bud (Gao et al., 2011). However, it is still unclear how Wnt5b protein can be transported through the tissue after lipid modification by Porcupine-mediated palmitoylation (Kurayoshi et al., 2007). Wg/Wnt ligands can be loaded on exovesicles (EVs) and activate the Wnt signaling cascade in the target cell (Gross et al., 2012). Indeed, recent reports suggest that the Wnt5b protein can be loaded on EVs of macrophages for long-distance transport to activate the invasion of breast cancer cells (Menck et al., 2013). In mice, Wnt5b is produced by hindbrain cells of the *Choroid Plexus* and associates with EVs, here lipoprotein particles, in the cerebrospinal fluid to orchestrate cerebellar morphogenesis (Kaiser et al., 2019). Finally, Wnt5b-associated exosomes also promote cancer cell migration and proliferation (Harada et al., 2017). However, it is unclear how Wnt5b can act in a paracrine way in a densely packed embryonic tissue like the zebrafish gastrula. The zebrafish glypican Gpc4 (*knypek*) has been suggested to control cell polarity during C&E in zebrafish (Topczewski et al., 2001). Recently, Gpc4 has been shown to be localized on signaling filopodia in zebrafish to aid Wnt/PCP signaling (Hu et al., 2021). Here, we show that Wnt5b can be loaded on cytonemes to facilitate paracrine signaling. These signaling protrusions can promote the exchange of lipophilic and membrane-tethered signaling components in zebrafish. Similarly, the lipid-modified Wnt proteins Wnt8a and Wnt3 are disseminated by cytonemes in zebrafish and gastric cancer cells, respectively (Routledge et al., 2022; Stanganello et al., 2015). Furthermore, we find that the cognate Wnt5b receptor Ror2 is localized on cytonemes. In support of our data, the Wnt receptor Fz in Drosophila and Fzd7 in chicks are also located on cytonemes to retrogradely transport the active Wnt receptors (Huang and Kornberg, 2015; Sagar et al., 2015). As an expansion to the cytoneme transport concept (Zhang and Scholpp, 2019), we show that Wnt5b can bind to Ror2 already in the producing cell and is then anterogradely transported along cytonemes to the receiving cells.

Intramolecular sensors based on fluorescence resonance energy transfer (FRET) have been employed for the analysis of conformational changes of several Wnt signaling components such as Fzd5 and Fzd6 (Kozielewicz et al., 2020; Wright et al., 2018) or Dishevelled (Harnoš et al., 2019) when exposed to Wnt5a. We used a refined procedure to quantify the ligand-receptor integrity during cytonemal transport in the living zebrafish embryo by FLIM-FRET. We could show the Wnt5b-GFP/mCherry-Ror2 complexes display a similar FRET efficiency and maintain the ligand-receptor distance in producing cells, on cytonemes, and in the receiving cells. These results indicate that the integrity of the complex is preserved during transport. Previously, ligand-receptor affinities have also been quantified in the Wnt signaling network. For example, an *in vitro* binding assay based on biolayer interferometry has suggested a binding affinity of Wnt5b to Fzd8 with a K_D_ of about 36nM (Bourhis et al., 2010). However, many known and unknown factors in the natural microenvironment may also influence the binding affinities of ligands to their receptors. Therefore, we used *in vivo* FCCS to precisely quantify the strength of receptor-ligand interactions in the living zebrafish. In the investigations presented here, we have measured *K*_*D*_ values of about 450nM for Wnt5b and Ror2. Similar to our analysis, FCCS analysis in Xenopus suggests that Ror2 can cross-correlate with Wnt5b; however, a K_D_ was not determined (Wallkamm et al., 2014). Instead, the analysis indicated that Wnt5b and Ror2 form large ligand-receptor complexes, which diffuse laterally in the membrane and are internalized to activate the pathway. Here we show that the binding affinity of Wnt5b to Ror2 is not altered during the complex transport from the producing to the receiving cell, indicating high structural integrity of the ligand-receptor complex.

After transport, we observe active signaling of the transferred Wnt5b/Ror2 complexes in the target cells. Supporting our finding, the dissemination of activated receptors on EVs has been observed in some disease contexts. For example, the cytokine receptor CCR5, the co-receptor for macrophage-tropic human deficiency virus (HIV)-1, can be released on EVs to CCR5 negative cells to enable infection of these cells (Mack et al., 2000). Furthermore, EGFRvIII in human glioma cells can be distributed by an EV-based transfer to activate other membrane-associated oncogenic tyrosine kinases such as Her-2, cKIT, or MET (Al-Nedawi et al., 2008). Such a mechanism could be operative in various human tumors to promote tumor progression, metastasis, and angiogenesis. However, cytonemal transfer of a receptor-tyrosine kinase-like Ror2 and their ligand in a physiological context has not been reported yet. In a parallel study by our group, we provide further evidence that in gastric cancer, ROR2, and the relevant ligand WNT5A are significantly correlated on cytoneme in human cancer-associated fibroblasts. We also identify these clusters in the cytoneme-receiving gastric tumor cells (Rogers et al., 2022). Moreover, cytoneme-dependent spreading of WNT5A/ROR2 complexes is critical for polarization and directed migration of the gastric cancer cells. Taken together, we suggest a conserved and essential role for this signaling mechanism in both vertebrate embryogenesis and disease.

Cell-cell communication can occur by several means, including chemical signaling. This form of communication involves the secretion of ligands as diffusible proteins or loaded on exosomes and cytonemes to engage with receptors of target cells at a distance. Here, we suggest a fresh cell-cell communication mechanism, namely the delivery of assembled and active ligand-receptor complexes. Such a mechanism increases the complexity by which cells may communicate. This communication can occur between directly adjacent cells; however, it can potentially also happen at a distance, as cytonemes have been observed with a length of hundreds of µm (Brunt et al., 2021). We suggest that cytonemes carrying the ligand and the receptor may be more effective in regulating the receiving cell behavior because the producing cell can target specific cells by cytonemes and precisely control the signaling strength within by adjusting the number of ligand-receptor pairs loaded on a cytoneme.

Furthermore, the availability of appropriate receptors in the target cells irrelevant. Overall, this work expands our knowledge about chemical signaling in embryogenesis.

During the development of all organisms, one of the most basic principles involves one group of cells changing the behavior of another group of adjacent cells. Generally, the principle of the “ability to react” formulated by Spemann at the beginning of the last century is based on multiple levels, including availability of signal transduction components, fitting set of transcription factor, and chromatin accessibility. However, the presence of the appropriate set of receptors at the top of the pathway remains the first essential prerequisite for signal activation, and by showing that cytonemes can deliver active ligand-receptor complexes, a revision of this long-standing concept is needed.

## Acknowledgments

Research in the Scholpp lab, including L.B., is supported by the BBSRC (Research Grant, BB/S016295/1 and an Equipment grant, BB/R013764/1) and by the Living Systems Institute, University of Exeter. C.Z. is supported by a Chinese Scholarship Council (CSC) studentship. S.R. is supported by the MRC (MR/S007970/1). We would like to thank Zhongxiang Jiang (Leica Microsystems) for his technical support in establishing *in vivo* FCCS and FLIM-FRET. We would further like to thank Thorsten Wohland (NUS Singapore), Gáspar Jékely, and Austin Smith (both LSI Exeter) for their critical comments on the manuscript.

## Author contributions

C.Z., S.R., and S.S. designed the experimental strategy. C.Z. performed and analyzed the experiments. L.B. performed the IF staining and generated the Ror2 CRISPR/Cas9 KO cell line. C.Z., S.R., and S.S. wrote the manuscript.

## STAR Methods

### RESOURCE AVAILABILITY

#### Lead contact

Further information and requests for resources and reagents should be directed to and fulfilled by the lead contact, Steffen Scholpp.

#### Materials availability

All the new plasmids we generated in this paper can be shared by the lead contact upon reasonable request.

#### Data and code availability

Microscopy data reported in this paper will be shared by the lead contact upon reasonable request.

Any information required to reanalyze the data reported in this paper is available from the lead contact upon request.

### EXPERIMENTAL MODEL AND SUBJECT DETAILS

#### Cell lines

PAC2 zebrafish fibroblasts were maintained at 28°C without CO2 in Leibovitz-15 media (Gibco, 11415056).

#### Zebrafish husbandry

Wild-type EZ9216B and *Tg (−6gsc: EGFP –CAAX)* (Smutny et al., 2017) zebrafish (*Danio rerio*) were maintained at 28°C on a 14 hr light/10 hr dark cycle (Brunt et al., 2021). All zebrafish husbandry and experimental procedures were followed and conducted under personal and project licenses granted by the UK Home Office under the United Kingdom Animals Scientific Procedures Act (ASPA) and following ethical policies, approved by the University of Exeter’s Animal Welfare and Ethical Review Body (AWERB). All the work with zebrafish was carried out before animals became capable of independent feeding, here at 5dpf or younger, pre-ASPA.

### METHOD DETAILS

#### Plasmids

Plasmids that were used for transfection, to generate mRNA for injecting into zebrafish embryos, and to generate probes for *in situ* hybridization experiments were listed as follows: zfGap43-GFP in pCS2+, xRor2^3i^ in pCS2+, xRor2-mCherry in pCS2+ and xRor2 in pCS2+ (Mattes et al., 2018). zfGap43 was amplified and cloned via BamHI and XbaI site to generate zfGap43-BFP in pCS2+. The open reading frame of xRor2 was amplified and ligated into pCS2+ mCherry using GeneArt™ Gibson Assembly HiFi Cloning Kit to make mCherry-xRor2. The open reading frame of zfRor2 was amplified and cloned via ClaI and XbaI sites into pCS2+ mCherry to generate zfRor2-mCherry. To generate zfRor2-ΔCRD-mCherry, the amino acids from 170 to 304 were deleted from zfRor2 ORF and cloned into pCS2+. The nanobody against GFP, the Sec-Vhh-mCh, was subcloned via ClaI and SnaBI site into pCS2+. The GPI-anchored mCherry was cloned into pCS2+ (Mem-mCherry) (Scholpp et al., 2009). The open reading frame of zfWnt5b was amplified and cloned via BamHI and XbaI site to generate zfWnt5b-GFP in pCS2+. Irsp53^4k^ (Brunt et al., 2021); Cdc42^T17N^ (Stanganello et al., 2015); pPBbsr-JNK KTR-mCherry (Miura et al., 2018)was a gift from Kazuhiro Aoki (Addgene plasmid # 115493), and we subcloned it into pCS2+ via ClaI and SnaBI sites; the antisense probe against *lef1* and *pax6a* were used as previously, cytosolic mCherry.

#### Transfection and CRISPR-Cas9 Knock Out

PAC2 zebrafish fibroblasts were maintained at 28°C without CO2 in Leibovitz-15 media (Gibco, 11415056). PAC2 cells were trypsinized and seeded on glass-bottom 35mm dishes for live imaging or on coverslips in 6-well plates for fixation. After 24hrs, cells were transfected with relevant plasmids using Fugene HD Transfection Reagent (Promega, E2312) and incubated at 28°C for 24hrs. Live cells were imaged on the Leica SP8 using the 63x water objective. To generate CRISPR Knock-out PAC2 cells, 50µM of Ror2 gRNA was generated from 100µM custom Ror2 crRNA (custom sequence: TACAACTGGAGCTCATCTGG, IDT DNA) and 100µM Alt-R CRISPR-Cas9 tracrRNA (IDT DNA, 1072532) and heated to 95°C for 5 mins and cooled to room temperature. 50µM of Ror2 gRNA was then incubated for 10-20 mins with 40µl nucleofector solution (Lonza P2 Primary cell 4D X kit L, V4XP-2024) and 20µM EnGen Cas9-NLS enzyme (NEB, #M0646T) to form the ribonucleoprotein (RNP) complex. 2×10^5^ PAC2 cells were centrifuged for 10mins at 1200rpm and washed in PBS, followed by 10mins centrifuge at 1200rpm. Cells were resuspended in nucleofector solution and combined with the RNP complex, PBS, and 100µM electroporator enhancer (1075915, IDT DNA) for 100µl total. Next, 100µl was transferred to a cuvette (Lonza P2 Primary cell 4D X kit L, V4XP-2024) and electroporated using a Lonza nucleofector. Next, 300µl pre-warmed Leibovitz-15 media was added to the cuvette and transferred to 2ml pre-warmed media in a 6-well plate and incubated at 28°C for 48hrs. For sequencing, DNA was extracted from cell pellets (GENEJET genomic DNA Purification kit, K0721, ThermoFisher Scientific), and the PCR product was amplified around the gRNA target site (Forward primer: CACACTTGAGACTTTGGGGGA; Reverse Primer: GGTGTAAAATCCTTACCTGC, Eurofins; PCRBIO, PCR Bio taq mix red, PB10.13-02). PCR products were sent for Sanger sequencing (Eurofins, TubeSeq Service).

#### Immunostaining

PAC2 zebrafish fibroblasts were seeded on coverslips in 6-well plates and transfected as above. After 24hrs, cells were fixed in 0.25% Mem-Fix (Rogers and Scholpp, 2020)(0.1M Sorensen’s Phosphate Buffer (pH 7.4), 4% formaldehyde, 0.25% glutaraldehyde) for 10mins at 4°C. Cells were washed 2x 5mins in Sorensen’s buffer and permeabilized in goat permeabilization buffer (0.1% Triton X-100, 5% goat serum, 0.2M glycine, 1× PBS) for 1hr at RT. Appropriate primary antibodies were used at 1:50 dilution in goat incubation buffer (0.1% Tween-20, 5% goat serum, 1× PBS). Primary antibodies used were: WNT5A-B, rabbit PolyAb, ProteinTech, 55184-1-AP; & ROR2 (D3B6F), rabbit mAb, Cell Signalling Technology, 88639S. 30µl of primary antibody in incubation buffer was placed on parafilm in a humidity chamber, and coverslips were placed cell-side down onto the buffer. Cells were incubated in primary antibody o/n at 4°C. Coverslips were placed cell-side up in 6-well plates and washed 6x 5mins in PBS. Appropriate secondary antibodies (Goat anti-rabbit IgG H&L Alexa Fluor 488, ab150077, Abcam) were prepared at 1:1000 dilution in goat incubation buffer. 30µl was placed on parafilm in a humidity chamber, and coverslips were placed cell-side down onto the buffer for 1hr at RT. Coverslips were then placed cell-side up in 6-well plates and washed 7x 10mins in PBS, then 1x in MilliQ and 1x in 1x PBS+0.05% Tween-20. Coverslips were mounted on slides using ProLong Diamond (Invitrogen) and left in the dark for 24hrs before imaging. Slides were imaged on the Leica SP8 using the 63x water objective.

#### Microinjection of mRNA

All the plasmids in this article were firstly linearised with corresponding NEW ENGLAND Biolabs, Inc (NEB) restriction enzymes. Then, capped sense mRNA was generated by in vitro transcription from linearized plasmids using Invitrogen mMessage mMachine SP6 Transcription kit. For different experiment purposes, zebrafish embryos at the 1-cell to 16-cell stage were injected with 1µl of different concentrations of mRNA. To generate clonal expression, Invitrogen Dextran, Fluorescein, and Biotin, 10000 MW (mini-Emerald) was co-injected with mRNA to the label producing or receiving cells.

#### Measurements of number and length of filopodia

Membrane-marked membrane protrusions were defined as filopodia as soon as they reached a length and width of 1µm. Zebrafish embryos were injected mRNA together with a membrane marker in 1 blastomere at the 8-cell to 16-cell stage. Injected embryos thus generated fluorescently labeled cell clones to visualize filopodia. Numbers of filopodia per cell were manually counted. Lengths of filopodia were measured from base to tips of protrusions in FIJI. At least 5 different embryos, and in each embryo, at least 10 isolated cells were measured.

#### Cell extension and convergence assay

Wild-type zebrafish embryos were injected with mRNA and mini-Emerald in 1 out of 8 to 16 cell blastomeres to generate a clonal expression. *Tg (−6gsc: EGFP –CAAX)* zebrafish embryos were injected with mRNA in 1 out of 8 to 16 cell blastomeres to generate a clonal expression. Then, at 9hpf, the embryos were mounted in a dorsal view for fluorescence imaging. First, the maximum width and length of a cell were measured in FIJI. In the meantime, the width of the notochord was also measured. Finally, the circularity was calculated to indicate the shape of the individual cell in the notochord. For the in situ hybridization analysis, the embryos were collected and fixed with 4 % PFA for further analysis after 30 hrs.

#### Image acquisition

Zebrafish embryos at 6hpf or 9-10 hpf were mounted in 0.7% low melting agarose in 35mm dishes. During the JNK inhibitor assay, mounted 6 hpf zebrafish embryos were treated with 40 μM JNK inhibitor (SP600125). Embryos were imaged with a Leica TCS SP8 confocal microscope, using a 63× dip-in objective. All the images were obtained from confocal z-stacks of living embryos.

#### *In situ* hybridisation

*Lef1* and *Pax6a* digoxigenin probes were generated using a Roche RNA labeling and detection kit and then purified through ProbeQuant G50 Micro Columns (Brunt et al., 2021).

Microinjected embryos with mRNA and mini-Emerald were collected at 5hpf and 30 hpf and fixed with 4% PFA. As previously described, in situ hybridization experiments were carried out (Brunt et al., 2021). In addition, we carried out double staining in these experiments to identify the producing cells. After the *Lef1* and *Pax6a* were properly stained, the embryos were fixed again and incubated at 70°C for an hour to deactivate the first probe. Later, we treated these embryos with antibody anti-FITC to bind the mini-Emerald. Finally, Fast-Red was used as a different color to label the clones. Double-stained embryos were kept in 70% glycerol for further analysis.

In situ embryos were imaged with an Olympus SZX16 stereomicroscope equipped with a DP71 digital camera. The images were taken using the Cell D imaging software. The embryos stained for lef1 were imaged with the animal pole up. Depending on the expression, we categorized the embryos into a regular expression, mild expression, and none expression to analyze the Wnt5b/Ror2 function. The embryos stained for pax6 expression were deyolked and flat-mounted in a dorsal view. The length of the forebrain and the width of the hindbrain were measured in FIJI. Based on the midline position and expression of pax6a in the hindbrain, we grouped them into three categories: normal phenotype, mild phenotype, and severe phenotype.

#### JNK reporter KTR-mCherry assay in vivo

The embryos were firstly injected at the one-cell stage with 250 ng/μl mRNA KTR-mCherry and kept in a 28°C incubator for 50mins. Later, we injected mRNA together with a membrane marker Gap43-GFP 1 out of 8 to 16 cell blastomeres to generate a clonal expression. When the embryos developed at 50% epiboly, the live sources were mounted and imaged. Cells that expressed GFP were considered producing cells. We drew a border to distinguish producing and receiving cells. Cells without nucleus KTR-mCherry expression were considered JNK signaling activation. We counted 5 cells distance away from the clone to analyze the paracrine JNK signaling activation. For each group, at least 6 different embryos were imaged.

#### Fluorescence cross-correlation spectroscopy (FCCS)

Before the FCCS measurements, we calibrated for the GFP channel (excited at 488nm) and the mCherry channel (excited at 587nm) to determine the effective volume (V_ef_). Therefore, the ATTO 488 and ATTO 565 dyes (Sigma Aldrich) with a known diffusion coefficient of 400 μm^2^/s (Kapusta; Müller et al., 2008) were used. The ATTO 488 was diluted into 3 nM and 6 nM to measure the auto-correlation in the GFP channel. The ATTO 565 was diluted into 4 nm and 8 nm to measure the auto-correlation in the mCherry channel. Finally, the effective volume for cross-correlation Vcc is determined in the following way (Schwille et al., 1997):

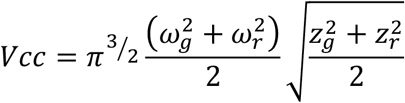

Embryos were injected in 1 out of 8 to 16 cell blastomeres with a low concentration of around 50 to 100 ng/μl. For FCCS measurements, the expression level has to be as low as possible. When the embryos were at 50% epiboly (6hpf), the live embryos were mounted in one of these cavities in 30 μl of 0.7% low-melting agarose and covered with a #1.5 coverslip. Tapes were used on both sides to stabilize the coverslip. The cross-correlation was measured by a Leica Sp8 FCS module equipped with FALCON single-molecule detection unit. Each measurement lasted 10s. The measurement procedure is illustrated in Suppl. Fig3.

For the measurement of the auto-correlation, the corresponding V_ef_ in each channel was applied (V_ef_ GFP 0.56 fl; V_ef_ mCherry 0.75fl), and the model of ‘Diffusion with Triplet’ was used for fitting. Vcc 0.65fl was applied for the cross-correlation fitting, and the ‘Pure Diffusion’ model was selected. All measurements were consistent with the calibration settings. The dissociation constant (Kd) was analyzed based on these fitting values for every measurement. Molecules in GFP focal volume is N1, and molecules in mCherry focal volume is N2. Molecules in the cross-correlation channel are Ncc, and the effective volume is Vcc. Based on these values from the fitting algorithm, the concentration in cross-correlation molecules was calculated as follows: NA is the Avogadro’s Constant.

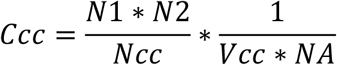

The concentration of molecules in the GFP channel is Cgreen and in mCherry is Cred. The Kd was calculated with the following equations.

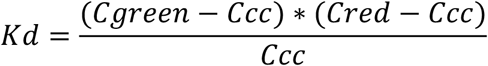

We excluded measurements in which the molecular concentration for each channel was over 2000nM.

#### Fluorescence Lifetime Imaging and Forster Resonance Energy Transfer (FLIM-FRET)

Embryos were injected in one blastomere with mRNA at the 8-cell to 16-cell stage. For FLIM-FRET experiments, we firstly injected the donor only to measure the donor’s lifetime. Then the donor and acceptor were co-injected to perform the FRET analysis. Injected 50% epiboly embryos were mounted in a plastic 30mm dish with 0.7% low-melting agarose. They were scanned with a Leica Sp8 FLIM module. The FLIM-FRET data were acquired by excitation at 488nm. Line repetition was set to 4 to collect enough photons.

The data were analyzed by using the LAS_X_Single Molecule Detection unit. The fitting model ‘Multi-Exponential Donor’ was selected for FRET analysis. The mean value of donor-only lifetime was applied to the unquenched donor lifetime to do the analysis. The Förster distance for the EGFP-mCherry pair is on the order of 52.4 Å (Akrap et al., 2010), which was also applied to the system. Different regions of interest (ROI) were selected and analyzed. The software calculated the mean lifetime, FLIM-FRET efficiency, and donor-acceptor distance.

#### Quantification and statistical analysis

Statistical analysis in these experiments was carried out using GraphPad Prism 9.0. Depending on different experiments, ordinary one-way ANOVA and Tukey’s multiple comparisons test, and two-way ANOVA together with Dunnett’s multiple comparisons test were used.

## Supplemental information

**Supplementary Figure 1.**
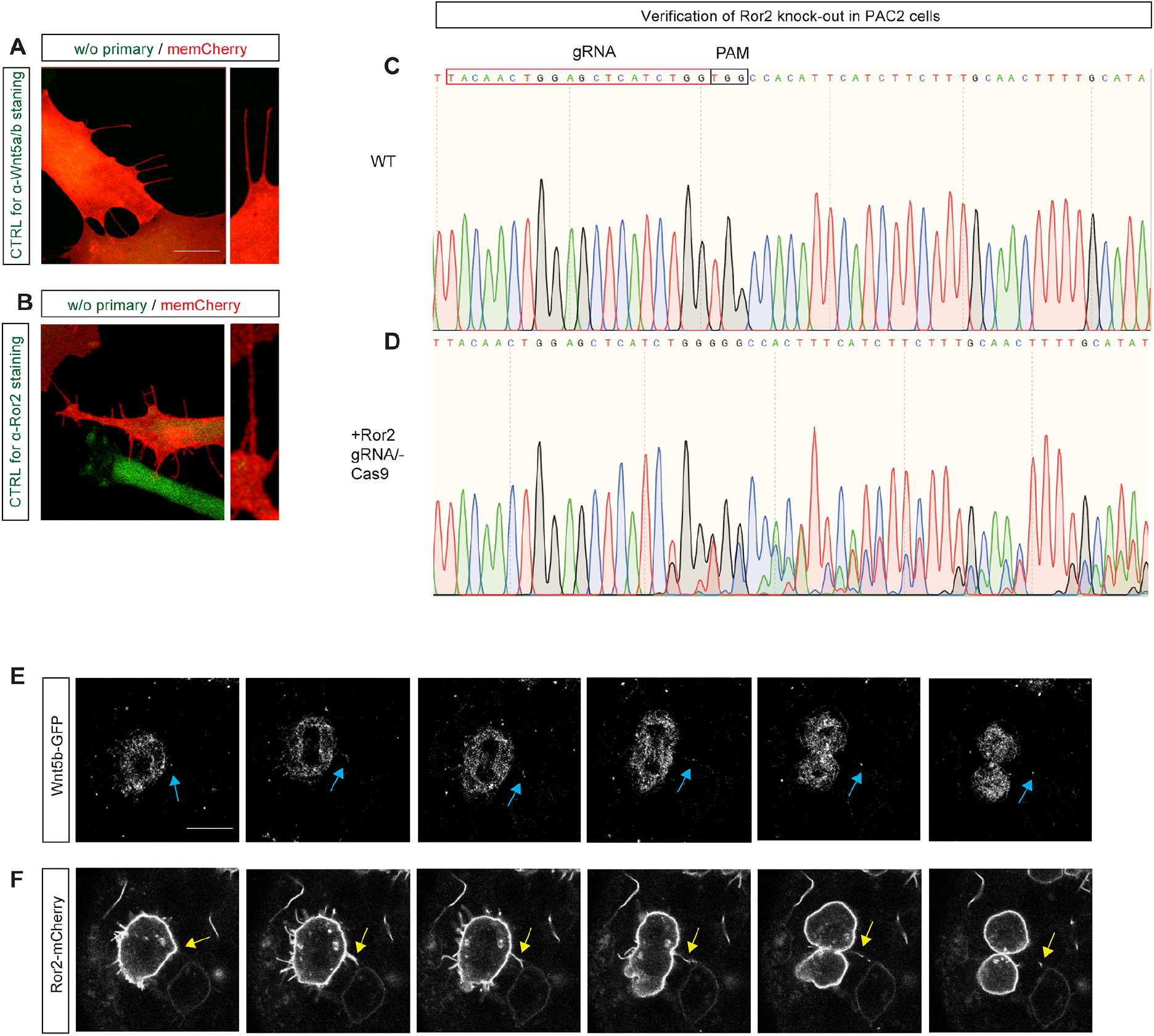
Wnt5b/Ror2 transport *in vitro* and *in vivo*. **A, B**. PAC2 zebrafish fibroblasts transfected with mem-mCherry and processed as described in Fig. 1A, B without primary antibodies. A high-magnification image indicates filopodia. Scale bar 10µm. **C, D**. Verification of CRISPR/Cas9-based knock-out of Ror2 by Sanger sequencing. Binding sites for gRNA and PAM are indicated. **E, F**. Time series of cytoneme-based transportation of Wnt5b-GFP and Ror2-mCherry in wild-type zebrafish embryos from 0’00’’ to 3’50’’. Embryos were injected with mRNA for Wnt5b-GFP and Ror2-mCherry at the 8-cell stage in one blastomere to generate local clones and imaged live at 6hpf. Yellow and blue arrows indicate Wnt5b-GFP and Ror2-mCherry localization on cytonemes, respectively. Scale bar 10µm.

**Supplementary Figure 2.**
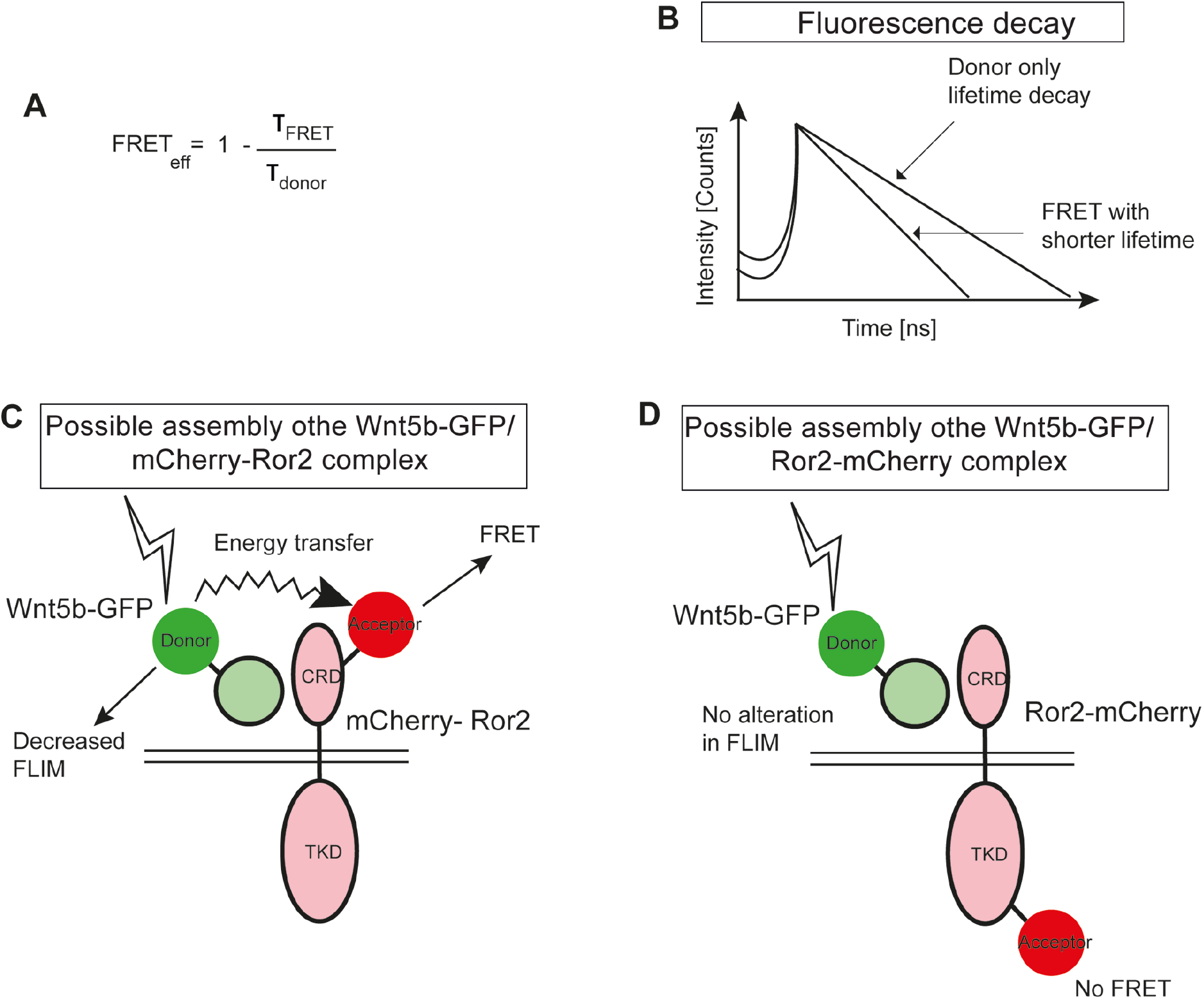
Principles of FLIM-FRET analysis of Wnt5b/Ror2 complexes. **A**. FRET efficiency, FRET_eff_, is defined as the fraction of photons that is transferred from a donor to an acceptor. τ_FRET_, a lifetime of the donor in the presence of the acceptor; τ_donor_, lifetime if the donor is without the acceptor. **B**. Bi-exponential fitting of the fluorescence decay demonstrates that the lifetime of a donor is decreased in the presence of a FRET acceptor. **C**. Close packaging in the Wnt5b-GFP/mCherry-Ror2 complex allows FRET to the acceptor and thus, reduces donor FLIM. **D**. A C-terminally fused Ror2-mCherry can be used as a negative control like the distance between donor and acceptor is too large, and the plasma membrane acts like as insulator.

**Supplementary Figure 3.**
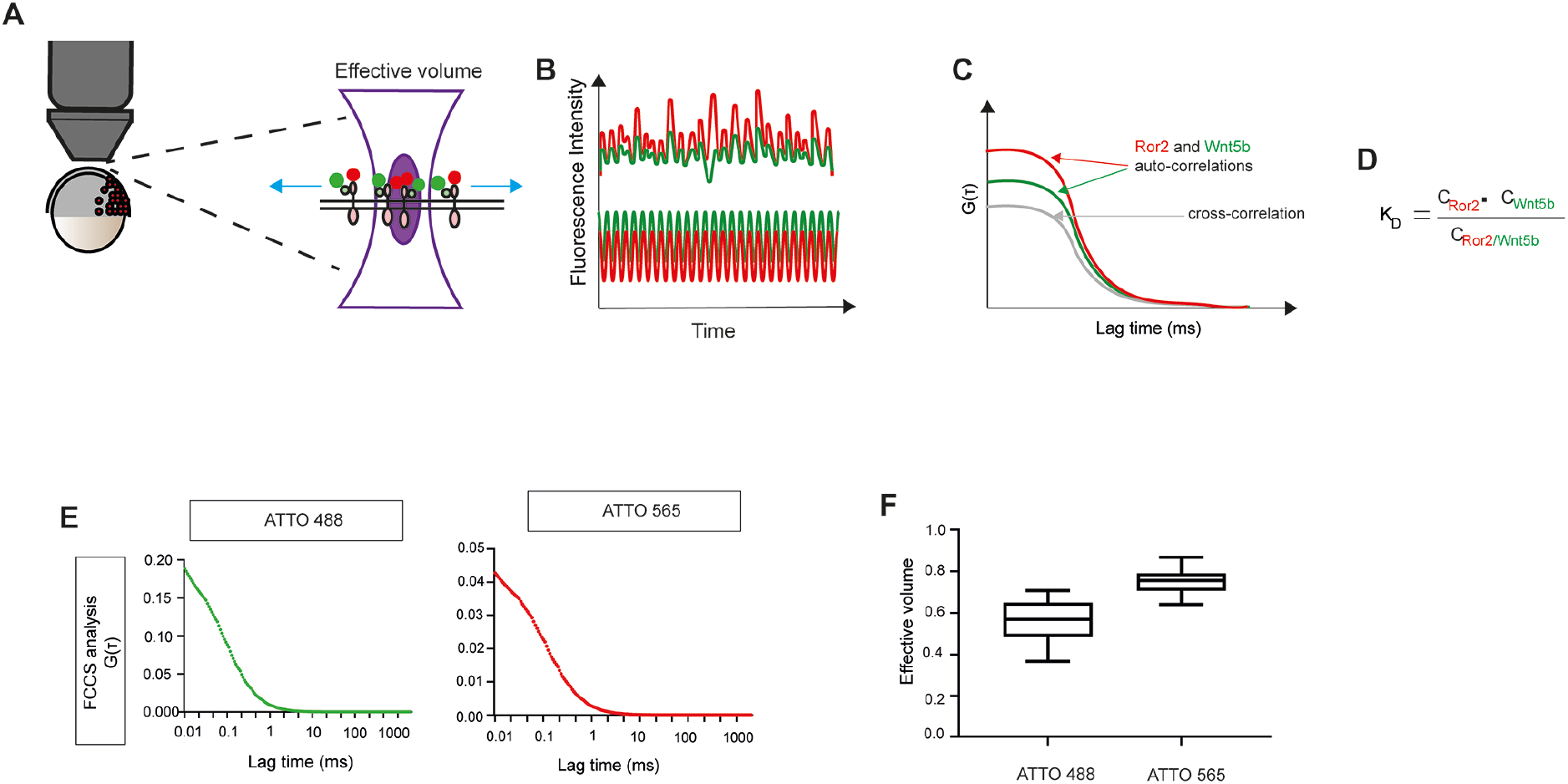
Principles of FCCS measurements of Wnt5b and Ror2. **A**. Schematic representation of the confocal detection volume positioned on the plasma membrane in a living zebrafish embryo to measure ligand-receptor interactions. **B**. Fluctuations in the detected fluorescence intensities with time. **C**. Auto- and cross-correlation analysis of fluorescence fluctuations over time. The Grey curve represents the cross-correlation between Wnt5b-GFP and Ror2-mCherry. **D**. The dissociation constant K_D_ is defined as the fraction of the concentration of the unbound ligands and unbound receptors to the concentration of the ligand-receptor complexes. **E**. Autocorrelation curves of ATTO 488 and ATTO 565 in solution to determine the FCCS volume. **F**. Determination of the FCS volume obtained from 10 different samples of labeled protein stock solutions.

**Supplementary Figure 4.**
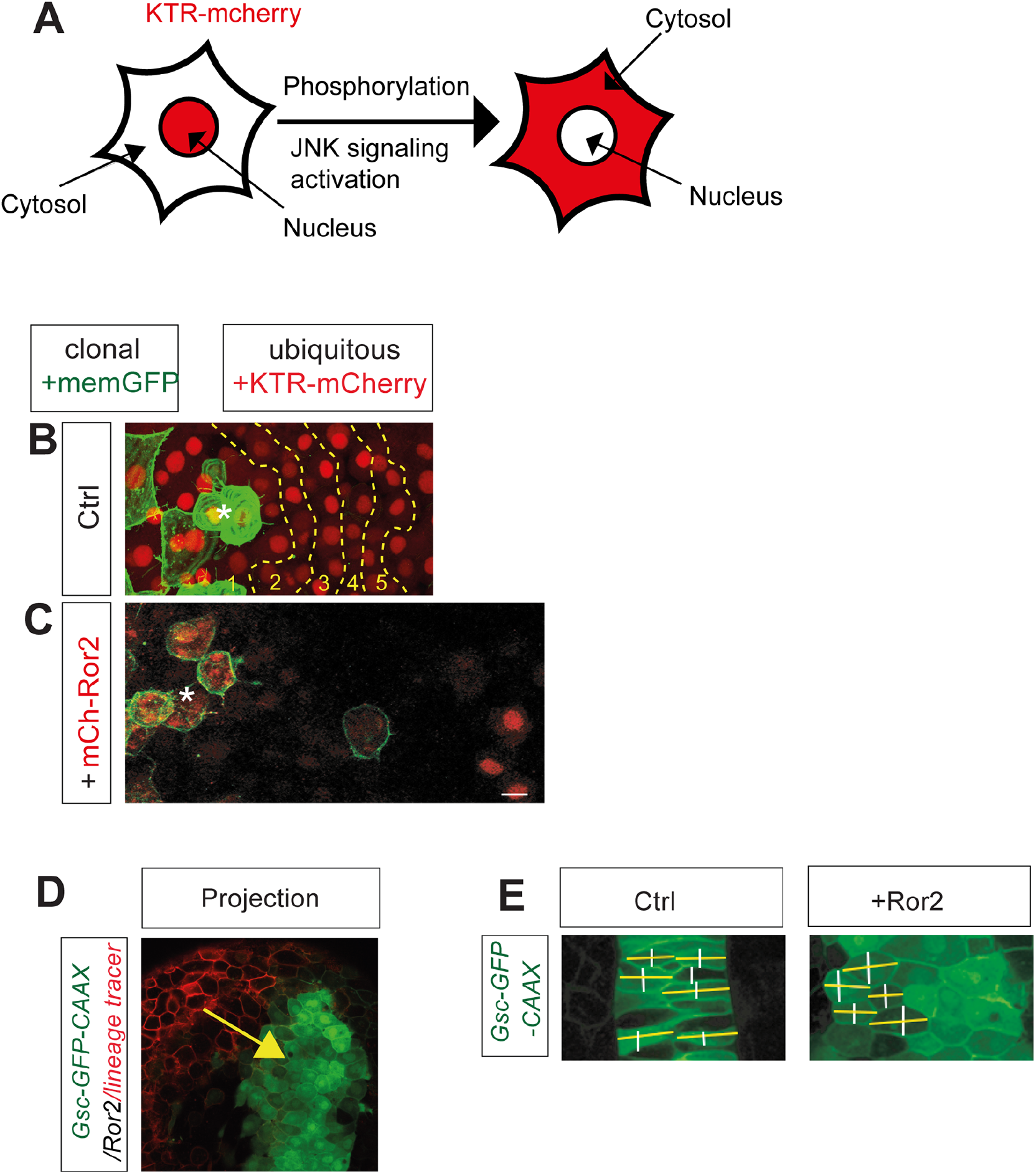
Principles of *in vivo* JNK reporter and JNK signal analysis. **A**. Principle of the JNK signaling KTR-reporter. **B**. Principle of measurements of cell rows around the Wnt5b/Ror2-expressing clones. **C**. N-terminal fusion of mCherry with Ror2 activates JNK signaling around the clone, similar to the C-terminal fusion in Fig. 5C. **D**. Circularity assay. Some example cells were measured to illustrate how the width of a cell in parallel to the body axis (white lines) or perpendicular to the body axis (yellow line) was measured in a WT embryo or an embryo with a Ror2 expressing clone at 24h. The ratio between the yellow line and the white line was calculated. A perfect circular cell has a circularity of 1.0, while below or above 1.0 indicates a noncircular, elongated shape.

